# Development of subunit selective substrates for *Trichomonas vaginalis* proteasome

**DOI:** 10.1101/2023.04.05.535794

**Authors:** Pavla Fajtova, Brianna M Hurysz, Yukiko Miyamoto, Mateus Serafim, Zhenze Jiang, Diego F. Trujillo, Lawrence Liu, Urvashi Somani, Jehad Almaliti, Samuel A. Myers, Conor R. Caffrey, William H. Gerwick, Christopher J Kirk, Evzen Boura, Lars Eckmann, Anthony J O’Donoghue

## Abstract

The protozoan parasite, *Trichomonas vaginalis* (Tv) causes trichomoniasis, the most common, non-viral, sexually transmitted infection in the world. Only two closely related drugs are approved for its treatment. The accelerating emergence of resistance to these drugs and lack of alternative treatment options poses an increasing threat to public health. There is an urgent need for novel effective anti-parasitic compounds. The proteasome is a critical enzyme for *T. vaginalis* survival and was validated as a drug target to treat trichomoniasis. However, to develop potent inhibitors of the *T. vaginalis* proteasome, it is essential that we understand which subunits should be targeted. Previously, we identified two fluorogenic substrates that were cleaved by *T. vaginalis* proteasome, however after isolating the enzyme complex and performing an in-depth substrate specificity study, we have now designed three fluorogenic reporter substrates that are each specific for one catalytic subunit. We screened a library of peptide epoxyketone inhibitors against the live parasite and evaluated which subunits are targeted by the top hits. Together we show that targeting of the β5 subunit of *T. vaginalis* is sufficient to kill the parasite, however, targeting of β5 plus either β1 or β2 results in improved potency.

## INTRODUCTION

*Trichomonas vaginalis* (Tv) is the parasite responsible for the most common non-viral sexually transmitted infection, trichomoniasis^1.2^. Trichomoniasis can cause a variety of symptoms including vaginal discharge, irritation, and itchiness, but since it is often asymptomatic it has historically been disregarded as a serious threat to human health and therefore diagnosis and treatments are understudied^3,4^. However, the rate of comorbidity with other sexually transmitted infections as well as the relationship with other serious health concerns including cancers, HIV acquisition, infertility, and adverse pregnancy outcomes points to a need for new diagnostics and therapies^3^. Due to a lack of screening for trichomoniasis, incidence estimates range anywhere from 156-276 million cases annually^4-6^.

The nitroimidazole drug family is the only FDA approved class of drugs for treatment of trichomoniasis, the two most common of which being metronidazole and tinidazole. Nitroimidazoles are prodrugs that are activated following nitro group reduction that occurs in certain microbes, leading to imidazole fragmentation and cytotoxicity^7^. While metronidazole and tinidazole are generally effective at treating trichomoniasis, side effects include nausea, abdominal pain, diarrhea, neurotoxicity, and optic neuropathy, while non-compliance, and reinfection are of concern^8^. Recent studies have also considered how oral metronidazole treatment impacts the gut microbiome. In a murine model, metronidazole was linked to a reduction of probiotic related bacteria communities and development of antibiotic resistant bacteria^9^. In healthy dogs, metronidazole decreased microbiome richness and reduced abundance of key bacteria that did not recover 4 weeks after discontinuing treatment^10^. Lastly, Tv strains with some resistance to metronidazole are becoming more prevalent, with some studies estimating resistance as high as 9.6%^8, 11-13^. The high incidence of Tv infection paired with poor diagnostics, correlation to other serious health impairments, growing resistance to drugs, and the deleterious effects of drugs on the gut microbiome, make the discovery of new trichomoniasis treatments imperative.

We previously validated the 20S proteasome as a drug target in Tv infections^14^. This enzyme is a high value target as few bacterial species have a proteasome, and therefore a proteasome inhibitor would have markedly fewer, if any, negative impacts on the gut microbiome than the current leading trichomoniasis treatment, metronidazole^15^. In eukaryotes, the 20S proteasome is responsible for proteolysis in the ubiquitin dependent protein degradation pathway. This process of degrading damaged or misfolded proteins is necessary for eukaryotes to maintain homeostasis. The proteasome has a hollow cylindrical structure consisting of two outer heptameric rings of α subunits and two inner heptameric rings of β subunits. The β rings contain three catalytic subunits, β1, β2 and β5, each with unique substrate specificity that varies by species. Inhibiting the human proteasome is a well-established strategy for treating cancers such as multiple myeloma^16^. However, these clinically approved drugs are likely to be too toxic for use in the treatment of parasitic diseases. The active site differences between the human and parasite can be exploited for developing inhibitors that specifically target the parasite proteasome and not the human proteasome^17-25^. We have previously developed proteasome inhibitors that are more than 2,000-fold more potent at killing *Plasmodium falciparum* cells than human cells^25^.

When initially validating the *T. vaginalis* 20S proteasome (Tv20S) as a drug target, we found little sequence conservation between the human and trichomonas catalytic subunits (28-52% sequence identity)^14^. Further, Tv20S was screened with a library of proteasome inhibitors derived from the marine natural product, carmaphycin B, and the leading compound, carmaphycin-17 (CP-17), was more effective at inhibiting Tv20S *in vitro* as well as *in vivo* using the murine model, *Tritrichomonas foetus*, than metronidazole. CP-17 also displayed the potential to overcome metronidazole resistance as it had similar effects preventing Tv growth in a metronidazole resistant strain. Lastly, CP-17 showed >5-fold greater potency to Tv20S proteasome than human proteasome and was less cytotoxic to human HeLa cells than anticancer proteasome inhibitors.

After validating the Tv20S proteasome as a potential drug target, we focused our attention on performing an in-depth characterization of the three catalytic subunits. The goal was to definitively identify each subunit using an activity-based probe and proteomics, to uncover the substrate specificity of each subunit, and to develop subunit-specific reporter substrates that can be used to elucidate the mechanism of action of proteasome inhibitors.

## RESULTS

### Proteasome purification and analysis

In this study, we developed a procedure to enrich for Tv20S from *T. vaginalis* protein extract using sequential steps of ammonium sulfate precipitation, size exclusion chromatography and anion exchange chromatography (Figure S1). At each step, catalytic activity was evaluated using a fluorogenic substrate, Suc-Leu-Leu-Val-Tyr-7-amino-4-methylcoumarin (Suc-LLVY-amc), which yielded a protein of ∼700 kDa (**Fig. 1A**). The molecular weight of Tv20S was found to be lower than the ∼750 kDa human constitutive 20S proteasome (c20s)^26^. To validate that this protein was Tv20S, we incubated it with a proteasome-specific activity-based probe, consisting of a tripeptide-vinyl sulfone inhibitor with a fluorescent tag^27^, and ran the enzyme-probe complex on a native gel. We clearly show that the probe binds to Tv20S in both the cell lysate and the enriched sample (**Fig. 1B**). These studies validate that Tv20S can be isolated with high purity from *T. vaginalis* lysate and that its molecular weight is lower than that of the human c20S. These studies correlate well with the calculated mass of the c20S (716.4 kDa) and Tv20S (697.3 KDa) using the protein sequence information (Supplementary File 1).

**Figure 1.**
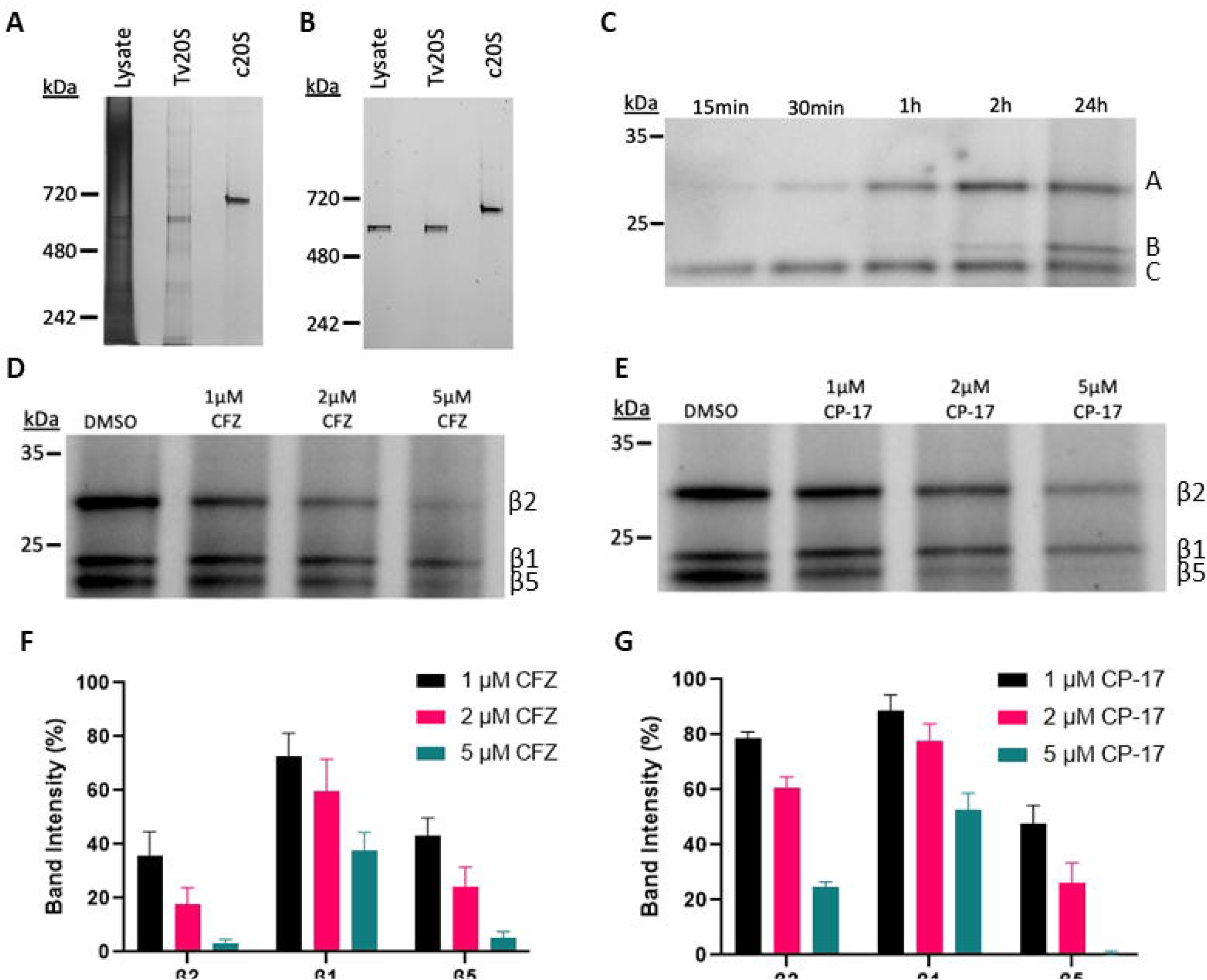
Gel based characterization of Tv20S. **A)** Tv lysate, purified Tv proteasome, and constitutive human proteasome were run on a NativePage gel, silver stained and imaged. **B)** The same protein samples were incubated with 2 μM of the fluorescent proteasome inhibitor probe, MV151, for 1hr and imaged at 470 nm excitation 530nm emission prior to silver stain. **C)** Tv lysate was incubated with 2 μM MV151 for up to 24hr and the catalytic subunits were imaged on a denaturing gel. **D)** Tv20S was preincubated with 1 μM to 5 μM of carfilzomib (CFZ) prior to labeling with MV151 and run on a denaturing gel. E) Tv20S was preincubated with 1 μM to 5 μM of CP-17 prior to labelling with MV151 and run on a denaturing gel. **F)** The bands from D were quantified with ImageJ software and the intensity of was graphed as a percentage of the bands in vechicle (DMSO) treated lanes **G)** The bands from E were quantified with ImageJ software and the intensity of was graphed as a percentage of the bands in vechicle (DMSO) treated lanes.

### Identifying catalytic subunits by gel band proteomics

To determine which subunit is labelled with MV151, we incubated the probe at increasing time-intervals from 15 min up to 24 h and then separated the individual subunits on a denaturing gel (**Fig 1C**). We show that the probe labelled a ∼25 and ∼30 kDa band within an hour and a third band with an intermediate molecular weight was labelled only after extensive incubation. This led us to the conclusion that MV151 labels different Tv20S subunits in a time dependent manner and requires 24 hours incubation to label all three bands. The three bands were excised from the gel and subjected to proteomic analysis. We determined that the upper band (Band A) corresponds to Uniprot protein A2F2T6. This protein has highest sequence homology to the human β2 subunit (Uniprot Q99436) and therefore we named the upper band Tv20S β2. Likewise, the middle (B) and lower (C) bands were determined to be Uniprot A2E7Z2 and A2DD57 that align with the human β1 (P28072) and β5 (P28074) subunits, respectively^14^.

Previously, carmaphycin analog 17 (CP-17) was found to be a potent inhibitor of *T. vaginalis*^14^. We show here that preincubation with 1 μM of CP-17 prior to MV151 showed a 55% reduction in β5 labelling (Band C), while β2 (Band A) and β1 decreased by 22% and 10%, respectively (**Fig. 1D**). At 5 μM the intensity of the β5 band decreased to undetectable levels while β2 and β1 decreased by 80% and 50%, respectively. When a similar assay was performed with the FDA approved inhibitor, carfilzomib, we observed that this compound binds to both β2 and β5 with >90% reduction in MV151 labelled at 5 μM. We conclude that CP-17 preferentially inhibits the β5 subunit of Tv20S while carfilzomib is a dual inhibitor.

### Uncovering the substrate specificity of *T. vaginalis* proteasome

To uncover the substrate specificity of Tv20S, β2 we incubated the enzyme with an equimolar mixture of 228 strategically designed synthetic peptides that have previously been validated as substrates for the human and Plasmodium proteasomes^21,28^. Quantification of the cleaved products by LC-MS/MS revealed that Tv20S cleaved 165 of 2,964 available peptide bonds after 20 h incubation (Supplementary File 2). We generated an iceLogo plot to illustrate the frequency of amino acids in each of the P4 to P4⍰ positions (**Fig. 2A**). In this profile, we revealed that hydrophobic amino acids are found most frequently in the P4, P2⍰ and P4⍰. At P3, there is a high frequency of hydrophobic residues in addition to positively charged R and H. At the P1 position, N, R and F are most commonly found, while peptides are never cleaved directly after P, G, T, K and Q. These studies also showed that 90% of the cleavage sites within the 14-mer peptides were located between the 4^th^ and 12^th^ amino acid, indicating that Tv20S strongly favored cleaving at sites that are distal from the N- and C-termini (**Fig. 2B**). This profile reveals that in *T. vaginalis*, the proteasome is an endopeptidase and is generally will not cleave proteins and peptides into single amino acids or di- and tri-peptides. Instead, it will degrade proteins into peptides that are greater than 4 amino acids in length.

**Figure 2:**
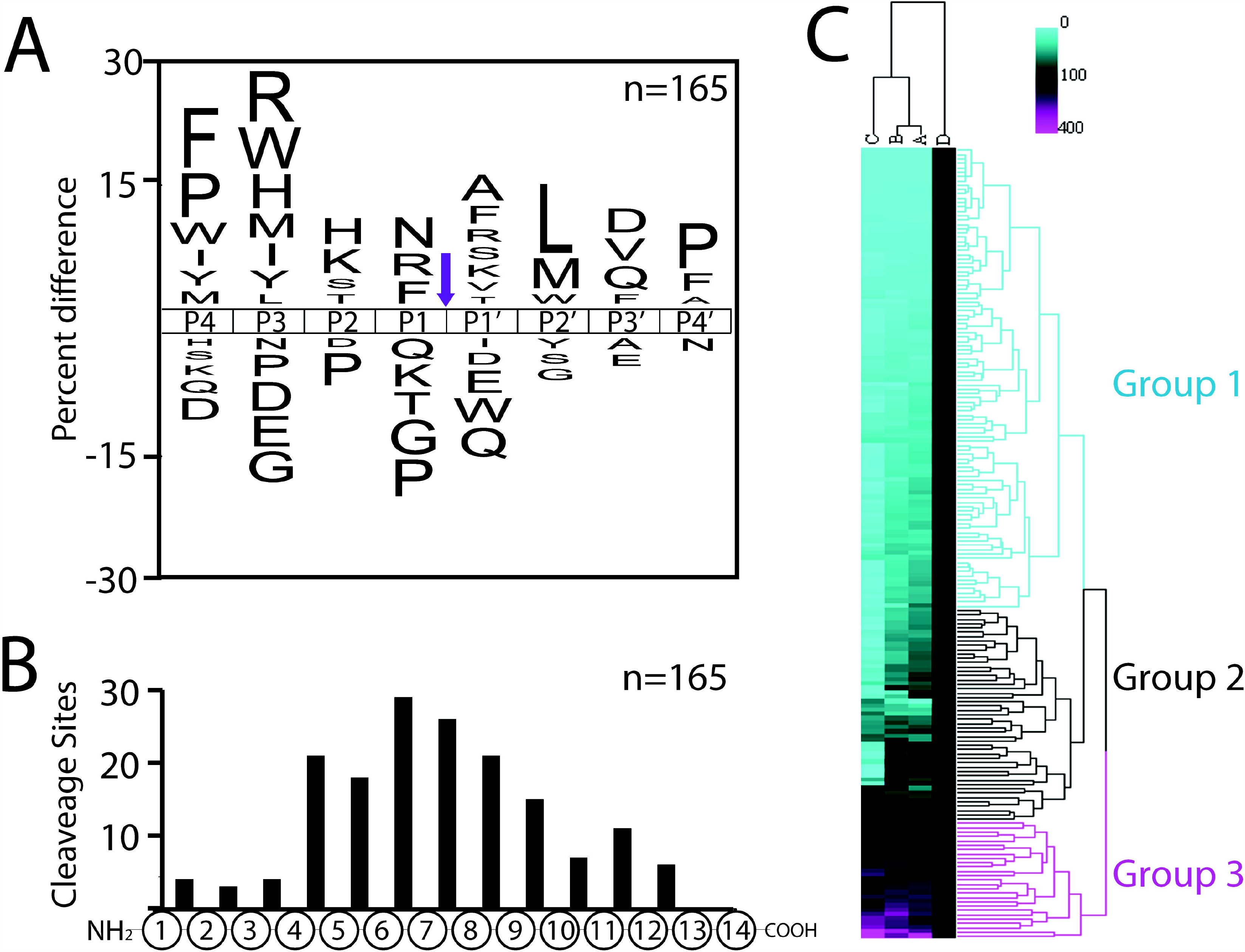
Characterization of peptides cleaved by Tv20S. **A)** IceLogo frequency plot of the peptides cleaved by Tv20S showing the amino acids that are increased (above mid-line) and decreased (below mid-line) in the P4 to P4⍰ positions. **B)** Distribution of cleavage sites within the 14-mer peptides. **C)** The intensities of cleavage products generated in the presence of 1 μM CP-17 (A), 10 μM CP-17 (B) and 10 μM CP-17+CFZ (C) were normalized to DMSO (D) and hierarchical clustering analysis was performed. Products clustered into three major groups.

Since Tv20S has six independent catalytic subunits, 2 each of the β1, β2 and β5 subunits, it is difficult to decipher which subunit is responsible for cleaving each peptide. The enzyme complex cannot be physically broken up into individual subunits for characterizing, and therefore we need to utilize inhibitors at defined concentrations to selectively inactivate one or more subunits. Therefore, we preincubated Tv20S with CP-17 and carfilzomib at concentrations that inactivate either β5 alone or β5 and β2 simultaneously. We then performed MSP-MS with these inhibitor-treated enzymes and compared the intensity of each cleaved product relative to the vehicle treated control (2.5% DMSO). From these normalized data, we performed hierarchical clustering analysis and found that three distinct groups of cleaved peptides were generated, which we attributed to each of the catalytic subunits of Tv20S (**Fig. 2C**).

### Development of a β5-specific Tv20S substrate

Group 1 had the largest number of cleaved peptides which corresponded to the β5 subunit. In this group, products did not form or significantly reduced in the presence of 1 μM of CP-17, conditions that we determined to only inhibit the β5 subunit. We generated an iceLogo of the 96 cleavage sites and revealed that this subunit preferentially cleaves peptides with hydrophobic amino acids in the P4, P1 and P4⍰ positions and mixed types of amino acids in other positions (**Fig. 3A**). Interestingly, the single most preferred amino acid was W at P3. The fluorogenic substrate that we have previously used for detecting activity of Tv20S β5 consists of the P4 to P1 sequence, LLVY. This substrate was developed for the human constitutive proteasome but only the Y residue in P1 appears to be preferred by Tv20S in the iceLogo. We chose a peptide in the library that contains W at P3 and a favorable hydrophobic residue (L) at P1. This peptide (AnTDRGWYL*AIQAV) was cleaved between L and A and we could quantify a time-dependent increase in the formation of the AnTDRGWYL product in the vehicle treated sample. The product was not formed in the presence of 1 μM of CP-17 therefore making it a β5 substrate (**Fig. 3B**). We synthesized the tetrapeptide substate, Ac-GWYL-amc, containing the P4 to P1 amino acids, flanked by an N-terminal acetylation group and a C-terminal fluorescent amc group. We evaluated the activity of Ac-GWYL-amc over a concentration range of 2 to 250 μM and calculated a K_m_ of 7.41 ± 1.73 μM. Under identical conditions, the K_m_ for the human β5 substrate, Suc-LLVY-amc was calculated to be 589.4 ± 134.7 μM. Therefore, we show that Ac-GWYL-amc is a superior substrate to Suc-LLVY-amc at all concentrations tested.

**Figure 3:**
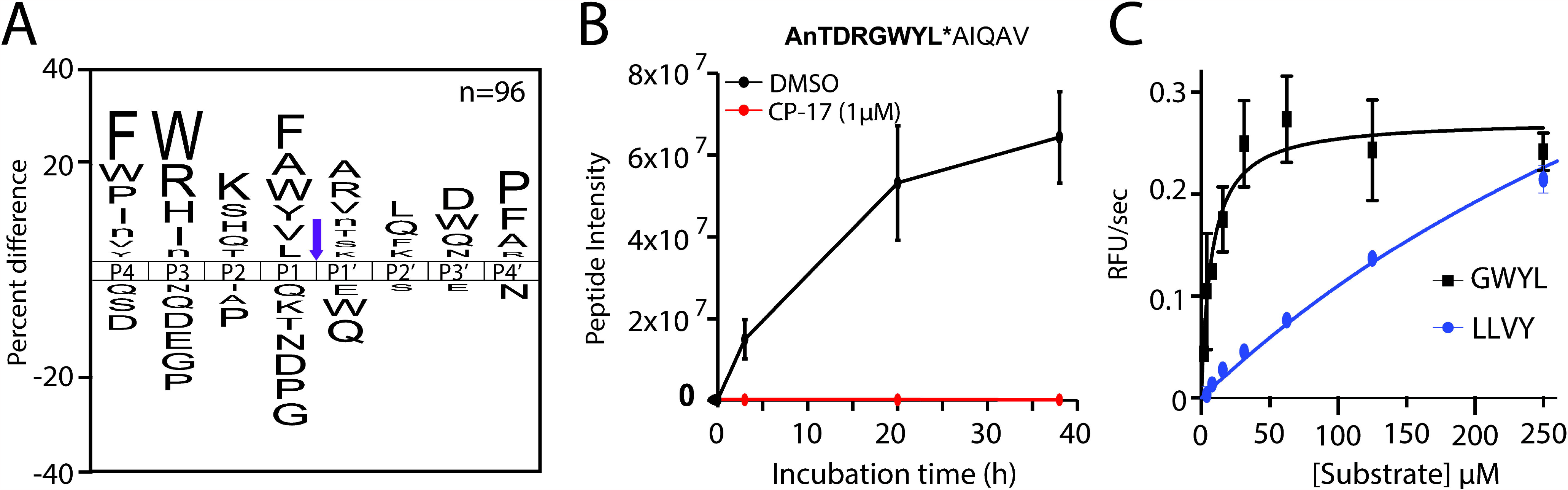
Design of a β5-specific substrate. **A)** IceLogo frequency plot showing the amino acids that are increased (above mid-line) and decreased (below mid-line) in the P4 to P4⍰ positions of the Tv20S β5 subunit. **B)** Example peptide that is cleaved by Tv20S and activity is inhibited by 1 μM CP-17. **C)** Michaelis Menten plot of Ac-GWYL-amc and Suc-LLVY-amc.

### Development of a β2-specific substrate

We next evaluated the peptides in Group 2 that are partially inhibited by 1 μM of CP-17 and have increased inhibition with 10 μM of CP-17 and a combination of CP-17 and carfilzomib. We anticipated that these cleaved peptides were generated from hydrolysis in the β2 subunit. We generated an iceLogo plot of the 48 peptides in that group and revealed that this subunit prefers cleaving peptides with hydrophobic amino acids in P4 and R in P3 and P1 (**Fig. 4A**). We chose HWAFRSR*YHGPLAH as a template for designing a β2 substrate as it closely matched the consensus sequence and synthesized Ac-FRSR-amc (**Fig. 4B**). We evaluated the activity of this substrate over a concentration range of 2 to 125 μM and calculated a K_m_ of 4.39 ± 1.0 μM. Under identical conditions, the K_m_ for the human β2 substrate, Z-LRR-amc was calculated to be 13.72 ± 2.36 μM (**Fig. 4C**). These studies confirm that Ac-FRSR-amc is cleaved at higher efficiency than Z-LRR-amc and is therefore an ideal reporter substrate for testing compounds that target the β2 subunit.

**Figure 4:**
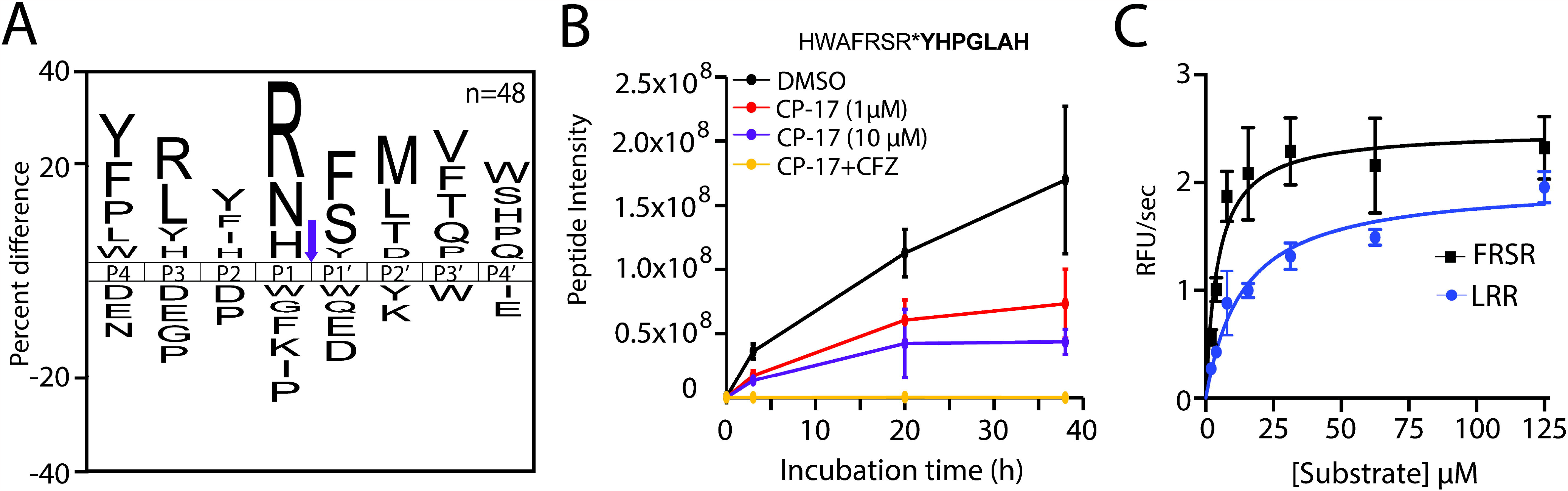
Design of a β2-specific substrate. **A)** IceLogo frequency plot showing the amino acids that are increased (above mid-line) and decreased (below mid-line) in the P4 to P4’ positions of the Tv20S β2 subunit. **B)** Example peptide that is cleaved by Tv20S, partially inhibited by 1 μM and 10 μM CP-17 and fully inhibited by 10 μM CP-17 and CFZ. The cleavage product in bold was quantified. **C)** Michaelis Menten plot of Ac-FRSR-amc and Suc-LLR-amc.

### Development of a β1-specific substrate

Finally, we evaluated peptides in Group 3 that were not inhibited by the combination of 10 μM CP-17 and 10 μM carfilzomib. In many cases the intensity of the cleaved products increased in the presence of these inhibitors and we predicted that these peptides were cleaved by the β1 subunit. We generated an iceLogo plot of the 27 cleavage sites in this group and revealed that most had either D, N or E in the P1 position (**Fig. 5A**). One of the most efficiently cleaved peptides in this group consisted of the sequence QYELPSRN*GWHHNP (**Fig. 5B**). The tetrapeptide sequence Ac-PSRN-amc was synthesized and cleavage of this substate was not inhibited by CP-17 or a combination of CP-17 and CFZ. Cleavage after N is uncharacteristic of the β1 specificity seen in other proteasomes^26^. Because of this, we hypothesized that cleavage of this substrate may be due to the presence of an abundant *T. vaginalis* protease, TvLEGU-1 that is known to hydrolyze peptides with P1-N. This protease may bind to Tv20S and therefore be co-purified with the enzyme. TvLEGU-1 cleaves the reporter substrate Z-AAN-amc^29^ and we predicted that it would be inhibited by RR11a, a broad-spectrum legumain inhibitor. We showed that Ac-PSRN-amc and Z-AAN-amc activities were inhibited by RR11a and therefore cleaved by TvLEGU-1 or a related enzyme (**Fig. 5C**). In search of a true β1 substrate, we chose EPLDLQKQRYFD*WL as a template because it was one of the most efficiently cleaved substrates containing a P1-D residue (**Fig. 5D**). We synthesized Ac-RYFD-amc and determined that it was not inhibited by RR11a confirming that it was not a legumain substrate. However, we did show that it was a β1 substrate as it was completely inhibited by bortezomib, a proteasome inhibitor that inactivates β1 activity in the human proteasome^30^ (**Fig. 4E**). Bortezomib did not inhibit activity with Ac-PSRN-amc and Z-AAN-amc thereby further confirming that those substrates are detecting an enzyme that is not Tv20S. We next compared the activity of Ac-RYFD-amc with the standard human β1 substrate, Z-LLE-amc. Surprisingly, Z-LLE-amc was more efficiently cleaved than Ac-RYFD-amc over the concentration range tested (**Fig. 5F**) even though the substrate profile revealed that Y at P3 is preferred over L and D at P1 is preferred over E. We predicted that Z-LLE-amc may be hydrolyzed by a subunit other than Tv20S β1. 10 μM of CP-17 alone or in combination with carfilzomib did not inhibit activity of the Ac-RYFD-AMC substrate, consistent with the MSP-MS data. In fact, activity increased by 160% or more as was also seen in the heatmap (**Fig. 2C**). However, using Z-LLE-amc, 1 μM of CP-17 partially inhibited activity while and a combination of CP-17 and carfilzomib completely inhibited activity (**Fig. 4F**). This cleavage profile is more similar to Tv20S β2 (**Fig 4**).

**Figure 5:**
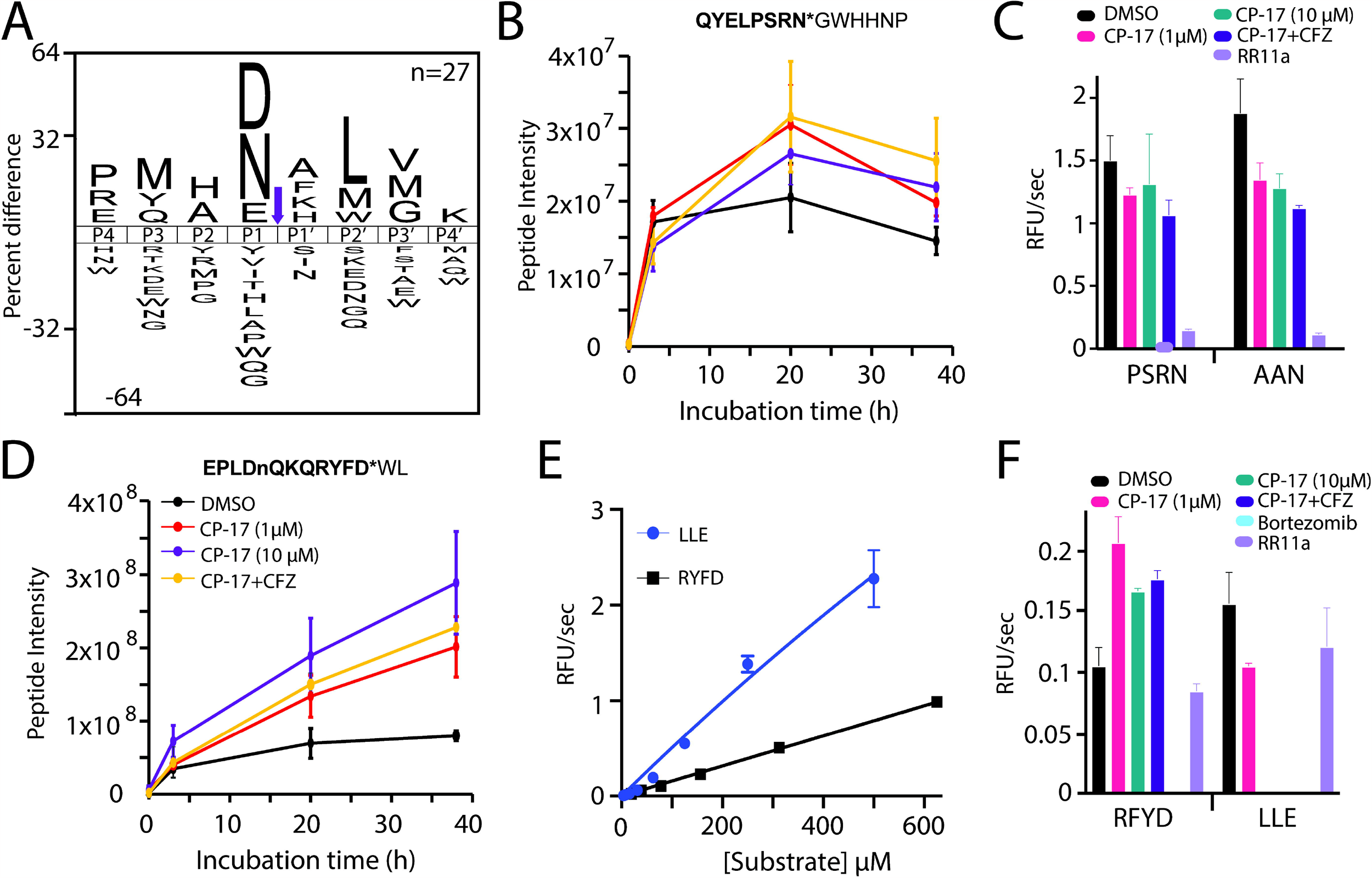
Design of a β1-specific substrate. **A)** IceLogo frequency plot showing the amino acids that are increased (above mid-line) and decreased (below mid-line) in the P4 to P4’ positions of the Tv20S Group 3. **B)** Example peptide product that is cleaved in the presence and absence of CP-17 and CFZ. **C)** Enzyme activity using Ac-PSRN-amc and the legumain substrate Z-AAC-amc. **D)** Example peptide product that is cleaved by Tv20S β1 where activity increases in the presence of CP-17 and CFZ. **E)** Michaelis Menten plot of Ac-RYFD-amc and Z-LLE-amc. **F)** Enzyme activity using Ac-RYFD-amc and Z-LLE-amc in the presence of proteasome and legumain inhibitors.

Taken together these data reveal that the MSP-MS was able to detect activity from all three catalytic subunits of Tv20S in addition to detecting legumain activity. Three subunit-specific fluorogenic reporter substrates were designed and shown to be superior to the commercially available substrates that were developed for human proteasome.

### Validation of substrates with previously discovered proteasome inhibitors

To validate the new rationally designed substrates for use in future Tv20S inhibitory screens, we generated dose-response curves with proteasome inhibitors such as leupeptin, ixazomib and carfilzomib, as well as our lead compound, CP-17^14^. In these assays, CP-17 was a potent β5 inhibitor (IC_50_ = 94.7 ± 18.9 nM) while carfilzomib inhibited β5 (IC_50_ = 78.4 ± 17.3 nM) and β2 (IC_50_ = 372.5 ± 92.1 nM) (**Fig. 6A, B**). For these compounds, we saw a concentration-dependent activation of β1. These subunit inhibition patterns were consistent with the decrease in MV151 labelling following preincubation with carfilzomib and CP-17 (**Fig. 1D, E**). Leupeptin has been shown to be an inhibitor of human β2^31^. Leupeptin also inhibited Tv20S β2 (IC_50_=3159 ± 654 nM) and activated β1. Finally, ixazomib was found to be a dual inhibitor of Tv20S β5 (IC_50_ = 34.1 ± 5.4 nM) and β1 (IC_50_ = 269.6 ± 97.1 nM) with activation of β2. Taken together these studies confirm that the three new subunit specific substrates are valuable tools for understanding the mechanism of action of different inhibitors.

**Figure 6.**
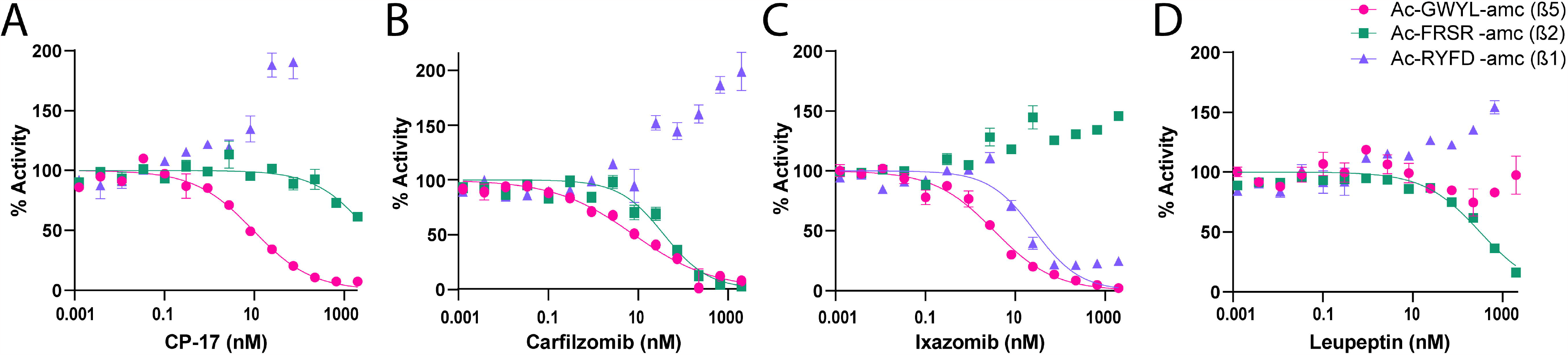
Use of new fluorogenic substrates with previously discovered proteasome inhibitors validates the substrates to be used for subunit specific proteasome inhibition screening. Dose-response curves for 0-20 μM of CP-17, carfilzomib, leupeptin, and ixazomib with activity being quantified using 30 μM Ac-GWYL-amc (β5), 30 μM Ac-FRSR-amc (β2) or 50 μM Ac-RYFD-amc (β1). Fluorescence was measured over 2h and the velocity was plotted to generate an IC_50_ curve.

### Screening of new inhibitors and characterizing subunit specific inhibition

To continue expanding upon our drug discovery work, we screened a library of proteasome inhibitors developed by Kezar Life Sciences, South San Francisco, CA, USA. These compounds are analogs of KZR-616, an immunoproteasome inhibitor in clinical trials for treatment of lupus nephritis^32^. The library contained 284 compounds that were initially screened in a microbial growth and survival assay, where ATP levels in Tv were used to measure parasite survival (**Fig. 7A**). After screening, two compounds were found with pEC50 > 6.5. KZR-1 had a pEC50 of 6.63 ± 0.13 (EC_50_ = 230 nM) and KZR-2 had a pEC50 of 6.56 ± 0.21 (EC_50_ = 280 nM). These compounds were then tested against purified Tv20S using the newly designed β1, β2 and β5 substrates (**Fig. 6B**). These data revealed that KZR-1 and KZR-2 only target Tv20S β5 with IC_50_ values of 136.0 ± 34.7 nM and 74.6 ± 24.3 nM, respectively. These compounds also increased β1 activity by 200% and thus we saw again that inhibition of β5 was correlated with an increase in β1 activity.

**Figure 7.**
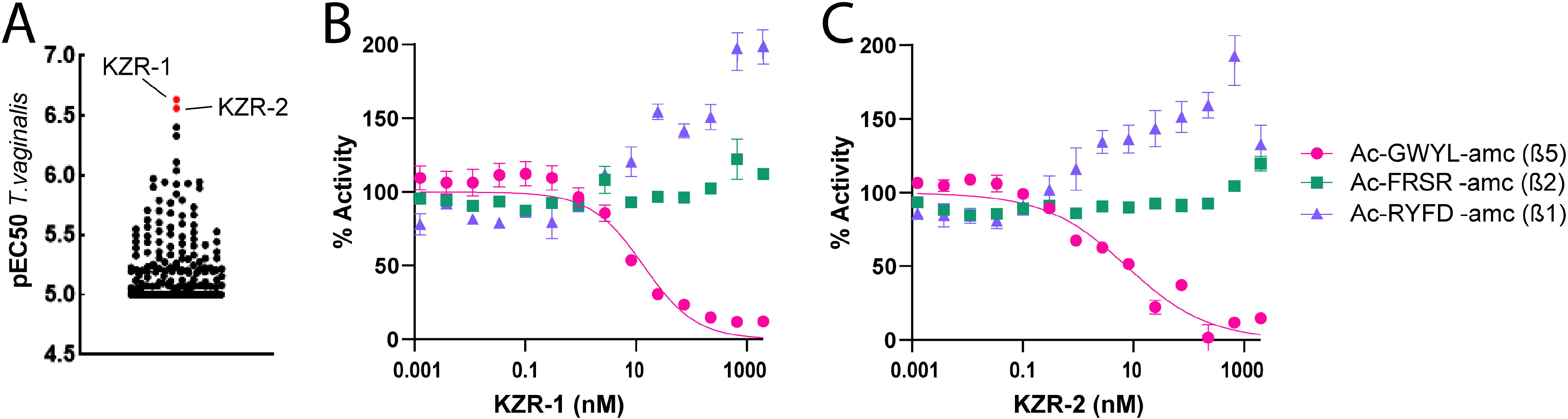
Screening of a new library of proteasome inhibitors provided two compounds that potently inhibit *T. vaginalis* growth with β5 subunit specificity. **A)** Anti-trichomonal activity of 284 proteasome inhibitors in Kezar Life Sciences library. Compounds were tested against *T. vaginalis* F1623 in 24h growth assays using ATP levels as readouts. Each data point represents the mean pEC50 for one compound. The top 2 most active compounds are highlighted in red. **B-C)** Dose-response curve for KZR-1 and KZR-2 ranging from 0.0125 nM - 20 mM were pre-incubated with 30 mM Ac-GWYL-amc (β5), 30 mM Ac-FRSR-amc (β2) or 50 mM Ac-RYFD-amc (β1) before Tv20S was added. Fluorescence was measured over 2 h and the velocity in RFU/s were plotted to generate an IC_50_ curve.

## DISCUSSION

Previously, we validated the Tv20S proteasome as a druggable target using a panel of peptide inhibitors with boronic acid or epoxyketone warheads^14^. We also screened a series of peptide-epoxyketone analogs and identified CP-17 as a hit compound that had highest selectivity. The present study advances our understanding of the catalytic function of the individual Tv20S subunits and provides us with improved tools for discovering new inhibitors. To do this, we first isolated Tv20S from *T. vaginalis* lysate by precipitating the proteasome using ammonium sulfate and then enriching the enzyme complex using size exclusion chromatography followed by anion exchange chromatography. The purity of Tv20S was higher than we previously reported from a two-step method^14^ and this allowed us to perform identify each of the catalytic subunits on a protein gel using proteomics. The fluorescent activity-based probe, MV151 binds to all three catalytic subunits, although at different rates. However, following a 16 h incubation, all three subunits are labelled, which facilitated excision of bands and identification of the protein subunits. Importantly, we showed that the covalent inhibitor CP-17 preferentially binds to the β5 subunit at low concentration (1 μM) while at 10-times higher concentration it inhibits β2. A second covalent inhibitor, carfilzomib, also targets β5 and β2 and therefore a combination of both CP-17 and CFZ was sufficient to chemically inactivate β5 and β2, thereby allowing us to biochemically characterize β1, the remaining catalytic subunit.

We have previously incubated the human constitutive 20S proteasome (c20S) and immune cell 20S proteasome (i20S) with a library of 228 tetradecapeptides and then quantified the cleavage products that are formed using LC-MS/MS^28^. The use of the peptide library combined with mass spectrometry is termed multiplex substrate profiling by mass spectrometry (MSP-MS). From these studies, i20S has an increased preference for cleavage of peptides on the C-terminal side of bulky hydrophobic amino acids such as tryptophan, whereas c20S has a greater preference for smaller and polar amino acids at that site. The tetrapeptide substrate Glu-Trp-Phe-Trp-7-amino-4-carbamoylmethylcoumarin (EWFW-acc) was subsequently developed to be an immunoproteasome-selective substrate while 5-methylisoxazolyl-Val-Gln-Ala-acc (iso-VQA-acc) was selective for c20S. These studies showed the value of MSP-MS for discovering differences between two related enzymes.

In this study, we used MSP-MS to uncover the substrate specificity of Tv20S β1, β2 and β5. However, the individual subunits cannot be physically separated from each other while still retaining catalytic function. Therefore, we used a combination of CP-17 and CFZ to selectively inactivate β5 or β5+β2 and then incubated the enzyme with the peptide library. These studies identified three groups of peptides which correlated with the three catalytic subunits. Peptides in group 1 were cleaved in the uninhibited reaction but were not cleaved in the presence of 1 μM of CP-17. Group 1 peptides are generated by Tv20S β5 and a fluorescent substrate Ac-GWYL-amc was developed as a reporter of β5 activity. Using the same approach, we also developed the β2 substrate, Ac-FRSR-amc using peptides cleaved in Group 2. Importantly, Ac-GWYL-amc and Ac-FRSR-amc were superior to the commercially available substrates, Suc-LLVY-amc (for β5) and Z-LRR-amc (for β2) that are commonly used for detecting human and microbial proteasome activities^33-35^.

In our studies of the peptides cleaved in Group 3, we found that many cleaved products were not inhibited by CP-17 and CFZ and some products were increased by up to 400% when compared to the untreated sample. In addition, a strong preference for cleaving peptides after Asn residues was usual, as the β1 subunits in proteasomes from other organisms generally cleave only after Glu, Asp and Leu^26^. We subsequently determined that the enzyme responsible for cleaving after Asn was a legumaim-like protease that co-purified with Tv20S. Co-purification of other proteases with proteasome has been reported previously^36^. For example, tripeptidyl peptidase II (TPPII) co-purified with the mouse proteasome from EL4 cells following ultra-centrifugation and anion exchange chromatography^37^. This led the authors to conclude that TPPII substitutes for some proteolytic activity of the proteasome. While TPPII is a very large protease with a MW of 138 kDa, the legumain-like proteases such as TvLEGU-1 are predicted to be ∼30 kDa^29^. This enzyme should not elute from the size exclusion column in the same fraction as the 700 kDa proteasome, leading us to hypothesize that it may bind to the proteasome. If so, the function of the legumain may be to provide the proteasome with additional specificity for cleaving after Asn residues. Our future studies will evaluate the role of this enzyme in Tv and determine if it is a direct binding partner of Tv20S.

When comparing the inhibition profile of the Tv20S β1 substrate, Ac-RYFD-amc with the commercial substrate Z-LLE-amc, we revealed that Z-LLE-amc is likely cleaved by a subunit other than β1. The MSP-MS data show that an increase in β1 activity occurs in the presence of CP-17 and CFZ exposure. This observation was then supported by studies using Ac-RYFD-amc where β1 activity increased by 160% in the presence of 10 μM CP-17. However, under the same conditions, Z-LLE-amc activity was eliminated by 10 μM CP-17. Therefore, Ac-RYFD-amc and Z-LLE-amc are not cleaved by the same subunit. Z-LLE-amc is likely to be hydrolyzed by both β1 and β5. While still a useful substrate to detect Tv20S activity in Tv extracts and to track proteasome activity in chromatography fractions during the purification procedure, Z-LLE-amc should not be used for directly evaluating Tv20S β1 activity.

Armed with three new subunit selective substrates, we validated their use in identifying the mechanism of action of different inhibitors. As expected, CP-17 was selective for β5 and inhibited β2 at higher concentrations while CFZ inhibited both β5 and β2. Ixazomib inhibited β5 and β1 but caused an increase in β2 while leupeptin inhibited β2 and increased activity of β1. These data reveal that the β1, β2 and β5 functions are dependent on each other. Based on the structure of the human proteasome, the β1 and β2 subunits are predicted to be adjacent on the β ring and therefore binding of an inhibitor to one subunit may directly affect substrate access to the adjacent subunit. It is unclear how inhibition of β5 modulates activity of β1 since they do not directly interact within the β ring. For future drug development efforts, it will be important to understand the cellular implications of inhibiting one or more subunits of the proteasome while simultaneously activating another subunit.

Finally, we tested our substrates using two new inhibitors that show good efficacy in *T. vaginalis* cultures. These compounds are analogs of KZR-616, an immunoproteasome selective inhibitor that is in clinical trials for treatment of lupus nephritis^32^. KZR-616 targets all three subunits of the human immunoproteasome, however its analogs inhibit only the β5 subunit of Tv20S with IC_50_ values of 136.0 ± 34.7 nM and 74.6 ± 24.3 nM. These compounds activate β1 in the nM range and β2 in the μM range and will become a starting point for development of a series of Tv20S selective inhibitors.

Taken together, our improved proteosome isolation methods and substrate specificity studies yielded three subunit selective substrates. These substrates are valuable tools for determining the mechanism of action of hit compounds from library screens. We have previously made potent human cathepsin B and Plasmodium proteasome inhibitors using substrate specificity data as a starting point for medicinal chemistry efforts^25, 38^. We will use the same approach here to develop compounds that are each selective for β1, β2 and β5. In addition, we will evaluate inhibitors that are derived for the LLE peptide found in Z-LLE-amc that may inhibit two subunits in parallel.

## ACKNOWLEDGEMENTS

The research was supported by NIH awards R01AI158612 and R21AI146387 to AJO and LE, R21AI133393 and R21AI171824 to AJO and CRC. PF received funding from the European Union’s Horizon 2020 research and innovation program under the Marie Skłodowska-Curie grant agreement No. 846688, ProTeCT. BMH and DFT were supported in part by the UCSD Graduate Training Program in Cellular and Molecular Pharmacology through an institutional training grant from the National Institute of General Medical Sciences, T32 GM007752. J.A. would like to acknowledge the St. Baldrick’s Foundation for the International Scholar award 2022–2025 and the deanship of scientific research at the University of Jordan for the scientific leave. MS would like to thank OpenEye Scientific for the academic licenses. MS was funded by CAPES foundation, grant numbers 88887.595578/2020-00 and 88887.684031/2022-00, and UFMG intramural funds.

## MATERIALS & METHODS

### *T. vaginalis* material

Culturing *T. vaginalis* F1623 parasites used in this study were grown in TYM (trypticase, yeast extract, maltose) Diamond’s medium supplemented with 180 mM ferrous ammonium sulfate at 37°C under anaerobic conditions.

### Purification of native Tv20S

Tv20S proteasome was purified from frozen pellets of Tv parasites using a modification of published procedure^14^. Frozen pellets of washed parasites in 50 mM HEPES, pH 7.5, 10 mM E-64, were thawed on ice and resuspended in 1 mL of lysis buffer containing 50 mM HEPES, pH 7.5, 10 mM E-64, 100 mM AEBSF, 1 mM pepstatin A and 1 mM DTT and three times sonicated on ice. The lysate was clarified by centrifugation at 30,000 g for 20 min. The 20S proteasome was precipitated with 30% and 60% ammonium sulfate and collected by centrifugation (30,000 g, 30 min). The pellet was resuspended in 50 mM HEPES, pH 7.5, 125 mM NaCl and purified by gel filtration using a Superose 6 Increase column (GE Healthcare). Fractions containing enzyme that cleaved Suc-LLVY-amc and were sensitive to 10 μM carfilzomib were pooled. The pooled fractions were applied to a 5 mL HiTrap DEAE FF column (GE Healthcare), and washed with 50 mM Tris-HCl, pH 7.5. 20S proteasome was eluted using a 0-1 M NaCl gradient in 50 mM Tris-HCl, pH 7.5. Fractions containing proteasome activity were pooled, buffer exchanged to 50 mM HEPES, pH 7.5, aliquoted and stored in -80°C.

### Protein Gels

Tv20S and c20S were diluted with 50 mM HEPES pH 7.5 then mixed with 2 μM MV151. Tv lysate was first diluted with 50 mM HEPES pH 7.5 and 10 μM E64 before 2 μM MV151 was added. After MV151 addition, samples were incubated at 37°C for up to 16 hours. For denaturing gels, samples were mixed with 4X Bolt LDS sample buffer (Thermo) containing DTT, boiled for 5 minutes and loaded into a Bolt 4-12% Bis-Tris (Thermo) or NuPAGE 12% Bis-Tris (Thermo) alongside PageRuler Plus pre-stained protein ladder (Thermo). Gels were run with 1X MES or 1X MOPS SDS buffers (Invitrogen) at 130V. For native gels, samples were mixed with 2X Novex Tris-glycine native sample buffer and loaded into NuPage Tris-glycine gels (Invitrogen) with NativeMark unstained protein standard (Thermo). Gels were run at 130V with Novex Tris-glycine running buffer (Invitrogen). All gels were imaged on Bio-Rad ChemiDoc XRS+ at 470nm excitation 530nm emission for MV151 probe visualization and imaged with white light for silver stain visualization. Silver stain was performed using Pierce™ Silver Stain Kit (Thermo). Tv20S purity was quantified by dividing the intensity of the MV151 band by the total intensity of the lane.

### Preparation of substrate and inhibitors

Substrates were custom synthesized by GenScript and dissolved in DMSO to 10 mM (Ac-GWYL-amc, Ac-FRSR-amc) or 50 mM (Ac-RYFD-amc). Substrates were stored at -20°C (Ac-GWYL-amc, Ac-FRSR-amc) or -80°C (Ac-RYFD-amc). Inhibitors were dissolved in DMSO to 10 μM and serially diluted on a black 384 well plate. Inhibitors and substrates were transferred using an acoustic liquid transfer system into low volume 384 well plates as needed.

### MSP-MS

Tv20S was pre-incubated with CP-17, carfilzomib (CFZ) or DMSO prior to mixing with an equimolar mixture of 228 14-mer peptides such that the final concentrations of each peptide was 0.5 μM and the final concentration of inhibitor was either 1 μM (CP-17) or 10 μM (CP-17 and CFZ). The exact concentration of Tv20S was unknown due to levels being below the detection limit for protein quantification kits. However, a 1:20 dilution of the enzyme stock was used. Reactions were incubated at rt and after 3, 20, and 38 h, 10 μL of this mixture was removed and the enzyme inactivated by mixing with 40 μL of 8M urea. A 0 h timepoint was generated by first mixing 6.6 μL of inhibitor/enzyme, incubating for 20 mins, then adding 3.3 μL of peptide mix. The reactions were performed in quadruplicate reaction tubes in assay buffer consisting of 50 mM HEPES pH 7.5, 10 μM E64, 100 μM AEBSF, 10 μM pepstatin, 1 mM DTT. The samples were then desalted and prepared for mass spectrometry as previously described^39^.

### Hierarchical clustering

Clustering of peptides sequences and their cleave patterns were assessed by hierarchical cluster analysis (HCA) with the ArrayTrack™ bioinformatics tool from the National Center for Toxicological Research (NCTR), Food and Drug Administration (FDA). ArrayTrack™ is a minimum information about a microarray experiment (MIAME) supportive tool, which provides linking between results with functional information and a given method (i.e., data biological relevance). HCA allows investigating grouping of samples or any data elements by their similarities profiles. Analysis was performed employing the dual cluster method, with Euclidean metric for distance measurements, and Ward’s linkage type. Lastly, we employed iFeatureOmega webserver (https://ifeatureomega.erc.monash.edu/iFeature2/index.html) to encode the peptides sequences’ data into boxplots visual representations, in order to facilitate visualization of statistical summaries for the generated features.

### Kinetic Tv20S Assays and Inhibitor Screens

Inhibition assays were performed by diluting Tv20S 1:20 in assay buffer (50 mM HEPES pH 7.5, 10 μM E64, 100 μM AEBSF, 10 μM pepstatin, 1 mM DTT) with the indicated substrate. For substrate validation as in **Fig. 4** and **Fig. 2B**, a final volume of 30 μL/well of 384 well black plates (Thermo) was used. Inhibitor IC_50_ curves were generated with 7.5 μL final volume on 384 well low volume black plates (Greiner bio-one). Fluorescence was measured for 2 h at excitation and emission on a Synergy HTX Multi-Mode Microplate Reader (BioTek, Winooski, VT).

### Antimicrobial Assays

The antitrichomonal activity against *T. vaginalis* F1623^40^ was examined as described previously^14^. Cells were grown at 37°C in TYM Diamond’s medium supplemented with 180⍰μM ferrous ammonium sulfate. Briefly, stocks of the test compounds were diluted in 0.1 μL of dimethyl sulfoxide (DMSO) to 2 mM, and 1:3 serial dilutions were made in a 384-well plate using Acoustic Liquid Dispenser (EDC Biosystems). 20 μL of trophozoites (10^3^/well) suspension in TYM were added to 384-well plates with diluted compounds, and cultures were incubated for 24⍰h at 37°C under anaerobic conditions (AnaeroPack-Anaero system). Growth and viability were determined with an ATP assay by adding BacTiter-Glo microbial cell viability assay reagent (Promega) and measuring ATP-dependent luminescence in a microplate reader. The 50% effective concentration (EC50) was derived from the concentration-response curves using GraphPad Prism 9 (GraphPad Software).

---

**Figure S1. Purification of Tv20S by A)** size exclusion chromatography on a Superose 6 Increase column. 0.5 mL of cytosolic fraction cleared by ammonium-sulfate precipitation was loaded onto column and equilibrated with 50 mM HEPES, 125 mM NaCl, pH 7.5. Fractions of 0.5 mL were collected. The activity was measured with the fluorogenic substrate Suc-Leu-Leu-Val-Tyr-AMC (red line) and with the proteasome inhibitor carfilzomib (black line), the active Tv20S contained fractions were pooled and loaded on **B)** a 5 mL HiTrap DEAE FF column. The column was washed with 50 mM Tris-HCl, pH 7.5. The elution was carried out by a linear gradient of 0-500 mM NaCl. Fractions of 1.5 mL were collected, and the proteasome activity was measured with the Suc-Leu-Leu-Val-Tyr-AMC substrate (red line). The active Tv20S fractions were pooled, concentrated and stored in -80°C.

