## Supplementary file for "Development of subunit selective substrates for *Trichomonas vaginalis* proteasome"

**Human 20S proteasome subunits**

**(propeptides are labeled yellow)**

```
>sp|O14818|PSA7_HUMAN Proteasome subunit alpha type-7 OS=Homo sapiens OX=9606 GN=PSMA7 PE=1 SV=1
MSYDRAITVFSPDGHLFQVEYAQEAVKKGSTAVGVRGRDIVVLGVEKKSVAKLQDERTVR
KICALDDNVCMAFAGLTADARIVINRARVECQSHRLTVEDPVTVEYITRYIASLKQRYTQ
SNGRRPFGISALIVGFDFDGTPRLYQTDPSGTYHAWKANAIGRGAKSVREFLEKNYTDEA
IETDDLTIKLVIKALLEVVQSGGKNIELAVMRRDQSLKILNPEEIEKYVAEIEKEKEENE
KKKQKKAS
>sp|P20618|PSB1_HUMAN Proteasome subunit beta type-1 OS=Homo sapiens OX=9606 GN=PSMB1 PE=1 SV=2
MLSSTAMYSAPGRDLGMEPHRAAGPLQLRFSPYVFNGGTILAIAGEDFAIVASDTRLSEG
FSIHTRDSPKCYKLTDKTVIGCSGFHGDCLTLTKIIEARLKMYKHSNNKAMTTGAIAAML
STILYSRRFFPYYVYNIIGGLDEEGKGAVYSFDPVGSYQRDSFKAGGSASAMLQPLLDNQ
VGFKNMQNVEHVPLSLDRAMRLVKDVFISAAERDVYTGDALRICIVTKEGIREETVSLRK
D
>sp|P25786|PSA1_HUMAN Proteasome subunit alpha type-1 OS=Homo sapiens OX=9606 GN=PSMA1 PE=1 SV=1
MFRNQYDNDVTVWSPQGRIHQIEYAMEAVKQGSATVGLKSKTHAVLVALKRAQSELAAHQ
KKILHVDNHIGISIAGLTADARLLCNFMRQECLDSRFVFDRPLPVSRLVSLIGSKTQIPT
QRYGRRPYGVGLLIAGYDDMGPHIFQTCPSANYFDCRAMSIGARSQSARTYLERHMSEFM
ECNLNELVKHGLRALRETLPAEQDLTTKNVSIGIVGKDLEFTIYDDDDVSPFLEGLEERP
QRKAQPAQPADEPAEKADEPMEH
>sp|P25787|PSA2_HUMAN Proteasome subunit alpha type-2 OS=Homo sapiens OX=9606 GN=PSMA2 PE=1 SV=2
MAERGYSFSLTTFSPSGKLVQIEYALAAVAGGAPSVGIKAANGVVLATEKKQKSILYDER
SVHKVEPITKHIGLVYSGMGPDYRVLVHRARKLAQQYYLVYQEPIPTAQLVQRVASVMQE
YTQSGGVRPFGVSLLICGWNEGRPYLFQSDPSGAYFAWKATAMGKNYVNGKTFLEKRYNE
DLELEDAIHTAILTLKESFEGQMTEDNIEVGICNEAGFRRLTPTEVKDYLAAIA
>sp|P25788|PSA3_HUMAN Proteasome subunit alpha type-3 OS=Homo sapiens OX=9606 GN=PSMA3 PE=1 SV=2
MSSIGTGYDLSASTFSPDGRVFQVEYAMKAVENSSTAIGIRCKDGVVFGVEKLVLSKLYE
EGSNKRLFNVDRHVGMAVAGLLADARSLADIAREEASNFRSNFGYNIPLKHLADRVAMYV
HAYTLYSAVRPFGCSFMLGSYSVNDGAQLYMIDPSGVSYGYWGCAIGKARQAAKTEIEKL
QMKEMTCRDIVKEVAKIIYIVHDEVKDKAFELELSWVGELTNGRHEIVPKDIREEAEKYA
KESLKEEDESDDDNM
>sp|P25789|PSA4_HUMAN Proteasome subunit alpha type-4 OS=Homo sapiens OX=9606 GN=PSMA4 PE=1 SV=1
MSRRYDSRTTIFSPEGRLYQVEYAMEAIGHAGTCLGILANDGVLLAAERRNIHKLLDEVF
FSEKIYKLNEDMACSVAGITSDANVLTNELRLIAQRYLLQYQEPIPCEQLVTALCDIKQA
YTQFGGKRPFGVSLLYIGWDKHYGFQLYQSDPSGNYGGWKATCIGNNSAAAVSMLKQDYK
EGEMTLKSALALAIKVLNKTMDVSKLSAEKVEIATLTRENGKTVIRVLKQKEVEQLIKKH
EEEEAKAEREKKEKEQKEKDK
>sp|P28066|PSA5_HUMAN Proteasome subunit alpha type-5 OS=Homo sapiens OX=9606 GN=PSMA5 PE=1 SV=3
MFLTRSEYDRGVNTFSPEGRLFQVEYAIEAIKLGSTAIGIQTSEGVCLAVEKRITSPLME
PSSIEKIVEIDAHIGCAMSGLIADAKTLIDKARVETQNHWFTYNETMTVESVTQAVSNLA
LQFGEEDADPGAMSRPFGVALLFGGVDEKGPQLFHMDPSGTFVQCDARAIGSASEGAQSS
LQEVYHKSMTLKEAIKSSLIILKQVMEEKLNATNIELATVQPGQNFHMFTKEELEEVIKD
I
>sp|P28070|PSB4_HUMAN Proteasome subunit beta type-4 OS=Homo sapiens OX=9606 GN=PSMB4 PE=1 SV=4
MEAFLGSRSGLWAGGPAPGQFYRIPSTPDSFMDPASALYRGPITRTQNPMVTGTSVLGVK
FEGGVVIAADMLGSYGSLARFRNISRIMRVNNSTMLGASGDYADFQYLKQVLGQMVIDEE
LLGDGHSYSPRAIHSWLTRAMYSRRSKMNPLWNTMVIGGYADGESFLGYVDMLGVAYEAP
SLATGYGAYLAQPLLREVLEKQPVLSQTEARDLVERCMRVLYYRDARSYNRFQIATVTEK
GVEIEGPLSTETNWDIAHMISGFE
>sp|P28072|PSB6_HUMAN Proteasome subunit beta type-6 OS=Homo sapiens OX=9606 GN=PSMB6 PE=1 SV=4
MAATLLAARGAGPAPAWGPEAFTPDWESREVSTGTTIMAVQFDGGVVLGADSRTTTGSYI
ANRVTDKLTPIHDRIFCCRSGSAADTQAVADAVTYQLGFHSIELNEPPLVHTAASLFKEM
CYRYREDLMAGIIIAGWDPQEGGQVYSVPMGGMMVRQSFAIGGSGSSYIYGYVDATYREG
MTKEECLQFTANALALAMERDGSSGGVIRLAAIAESGVERQVLLGDQIPKFAVATLPPA
>sp|P28074|PSB5_HUMAN Proteasome subunit beta type-5 OS=Homo sapiens OX=9606 GN=PSMB5 PE=1 SV=3
MALASVLERPLPVNQRGFFGLGGRADLLDLGPGSLSDGLSLAAPGWGVPEEPGIEMLHGT
TTLAFKFRHGVIVAADSRATAGAYIASQTVKKVIEINPYLLGTMAGGAADCSFWERLLAR
QCRIYELRNKERISVAAASKLLANMVYQYKGMGLSMGTMICGWDKRGPGLYYVDSEGNRI
SGATFSVGSGSVYAYGVMDRGYSYDLEVEQAYDLARRAIYQATYRDAYSGGAVNLYHVRE
DGWIRVSSDNVADLHEKYSGSTP
>sp|P49720|PSB3_HUMAN Proteasome subunit beta type-3 OS=Homo sapiens OX=9606 GN=PSMB3 PE=1 SV=2
MSIMSYNGGAVMAMKGKNCVAIAADRRFGIQAQMVTTDFQKIFPMGDRLYIGLAGLATDV
QTVAQRLKFRLNLYELKEGRQIKPYTLMSMVANLLYEKRFGPYYTEPVIAGLDPKTFKPF
ICSLDLIGCPMVTDDFVVSGTCAEQMYGMCESLWEPNMDPDHLFETISQAMLNAVDRDAV
SGMGVIVHIIEKDKITTRTLKARMD
>sp|P49721|PSB2_HUMAN Proteasome subunit beta type-2 OS=Homo sapiens OX=9606 GN=PSMB2 PE=1 SV=1
MEYLIGIQGPDYVLVASDRVAASNIVQMKDDHDKMFKMSEKILLLCVGEAGDTVQFAEYI
QKNVQLYKMRNGYELSPTAAANFTRRNLADCLRSRTPYHVNLLLAGYDEHEGPALYYMDY
LAALAKAPFAAHGYGAFLTLSILDRYYTPTISRERAVELLRKCLEELQKRFILNLPTFSV
RIIDKNGIHDLDNISFPKQGS
>sp|P60900|PSA6_HUMAN Proteasome subunit alpha type-6 OS=Homo sapiens OX=9606 GN=PSMA6 PE=1 SV=1
MSRGSSAGFDRHITIFSPEGRLYQVEYAFKAINQGGLTSVAVRGKDCAVIVTQKKVPDKL
LDSSTVTHLFKITENIGCVMTGMTADSRSQVQRARYEAANWKYKYGYEIPVDMLCKRIAD
ISQVYTQNAEMRPLGCCMILIGIDEEQGPQVYKCDPAGYYCGFKATAAGVKQTESTSFLE
KKVKKKFDWTFEQTVETAITCLSTVLSIDFKPSEIEVGVVTVENPKFRILTEAEIDAHLV
ALAERD
>sp|Q99436|PSB7_HUMAN Proteasome subunit beta type-7 OS=Homo sapiens OX=9606 GN=PSMB7 PE=1 SV=1
MAAVSVYAPPVGGFSFDNCRRNAVLEADFAKRGYKLPKVRKTGTTIAGVVYKDGIVLGAD
TRATEGMVVADKNCSKIHFISPNIYCCGAGTAADTDMTTQLISSNLELHSLSTGRLPRVV
TANRMLKQMLFRYQGYIGAALVLGGVDVTGPHLYSIYPHGSTDKLPYVTMGSGSLAAMAV
FEDKFRPDMEEEEAKNLVSEAIAAGIFNDLGSGSNIDLCVISKNKLDFLRPYTVPNKKGT
RLGRYRCEKGTTAVLTEKITPLEIEVLEETVQTMDTS

Assembled human 20S proteasome calculated using Expasy (https://web.expasy.org/compute_pi/)
Theoretical pI/Mw: 5.98 / 358217.50 = 716,435 Da


Trichomonas vaginalis 20S proteasome subunits
Predicted propeptides are labeled yellow

>TVAG_152400  | Trichomonas vaginalis G3 | Family T1, proteasome beta subunit, threonine peptidase | protein | length=224 beta6
MEGEFRENKKGQWSPYEMHGGTAIGICGDDYVVIGADTRLSVDYSIDSRHKARIFKMNSN
CMISATGFDGDIDAFITRMRSILLNYENQHFHEMSVESVARCVSNTLYSKRFFPYYINIL
VGGINSEGKGKLYGYDPVGTIEDLHYDSNGSGSSLAAPLLDSAFGTIHHNTRPFPAVSLQ
DAKNIVRDAICSVTERDIYTGDALQLCVFTKDGFAQEEFPLPRH
>TVAG_267300  | Trichomonas vaginalis G3 | Family T1, proteasome alpha subunit, threonine peptidase | protein | length=241 alfa1
MSSGADRYLTVFSAEGRLWQVEYSFKAVKQAEVTAVAVKSKNAVCVAVQKKVSDKLIDPS
TVTHMYRITDNVGACLVGLPSDVNFIVMLLRSFANNFEYKQGFSIPVSILAQMLSERHQL
ESQLVYVRPSAVSAILFGLDGPSDSFALYKIEPSGYSNGFRAVACGVKEIEAMSALEKKM
EDFETPEATAEFTLSTLQTVCGVDFEAQDVEVSLLTRDNSKFSKLPNDKVNEILHAVAEK
D
>TVAG_185200  | Trichomonas vaginalis G3 | Family T1, proteasome alpha subunit, threonine peptidase | protein | length=240 alfa7
MSGAGSGYDFNPITFSPDGRQFQVEYATKAVEKDSLALGVKCKDGILLAAEKNLTSTLLT
PGGNPRIFWINDSIACATIGHRPDCYSIVEQSRNRAETFTSNFGIKITVPQLASEVSQQF
HLAHYYQAYRPFGCTVIFASYKDDALYAIEPSGAFYGYFASCFGKNSNLARAELQKTEWK
NITVREAVPEVARIIKSLHESQFKKWEIEMFWLCEETNGRPQKVPEDVFQSRFVNENPQN
>TVAG_286960  | Trichomonas vaginalis G3 | Family T1, proteasome beta subunit, threonine peptidase | protein | length=206 beta3
MSDISTYNGSCVLAMAGDHCVAIASDRRLGVNMLTVSKDFKRIFQINDRIYLGLAGLATD
VLTVREQLRFDVNLLELREERPIDPKKFMNLVKSTLYEKRFSPFFVTPVIAGLLPETNEP
YLAASDSIGAFAFPKDFAVAGTCEESLYGICESAWRPNMNPDELFECTAKCLIAAVERDS
ISGWGGIVYIITQDKVIIKEIKTRMD
>TVAG_013010  | Trichomonas vaginalis G3 | Family T1, proteasome beta subunit, threonine peptidase | protein | length=256 beta5
MQSLYLKPRDEVESEETTKALEPAHVPDPCQFVKNHISLSYTNEPGKMAAVHGTTTLSFI
YNGGIVVAVDSRATGGQFIFSQTVMKILPLAPNMIGTMAGGAADCQYWLRNLSRLIQLHK
FRYQQPLTVAAASKILVNELYRYKGYNLSIGSMICGYDNTGPHIFYIDNHGSRIAGKRFS
VGSGSTHAYGVLDTCYREDMTKEEACELGRRAIYHATYRDSGSGGRVSVVHITQNGVEWI
DKTDVFDMHDFSKTTF
>TVAG_340740  | Trichomonas vaginalis G3 | Family T1, proteasome alpha subunit, threonine peptidase | protein | length=235 alfa4
MSDYTRSITRFSPDGRLFQIDHAHAAVQRGTTVVATRSKDMIVIAVEKTAVAKLQDPHTF
SKICSLDKHVMCAFAGLHADARRLIQSGQRQCQSHRLTYEDPISIENIARYIATLQLKNT
QSGGARPYGVSTLICGFDDMTSQPHIYETLPSGTYAEWKARTIGRHDQTVMEYLEKHYKD
DMTDEEAQKLAIGALLEVVENGSKNLEVAYMKRGGTMEIMAEEVLDALIESTKAK
>TVAG_127250  | Trichomonas vaginalis G3 | Family T1, proteasome beta subunit, threonine peptidase | protein | length=216 beta1
MSEYQFPKESMGTTLLAIQCTDGVVMASDSRTSSGSFIPNRATNKITEIQPKIFAARCGN
AADTQFLARAVKNYLNALNITRENTDDSTILVASNVIRSLIVRYRQYLSAGVIVGGWDSA
GPQVYSIEVSGMAIKKKIASNGSGSTYIQAYIDQNYREDMTMEEATKFAIAAVTGAIIRD
GSSGGVVNIVQINADGAKRMTVRPAQQPFNYDIVKG
>TVAG_293540  | Trichomonas vaginalis G3 | Family T1, proteasome alpha subunit, threonine peptidase | protein | length=251 alfa5
MFNSGSEYDRNVNTFSPDGRLLQVEYAIEAVKLGSSAVAILCPEGVIFAVEKRLSSQLLI
ASSVEKVYAIDDHVGVVMAGLAADGRTMVEHMRVEAQNHRFSFDEPIGIKAVTQSVCDLA
LAFGEGRRKKGDGQMSRPFGTALLVAGIENGKCHLFHTDPSGTYTECRARAIGGGSEGAE
ALLRDLYKDGMTLHEAEDLALSTLRQVIQEKLNENNVEVACARVSTGKFEIYTSEQRQEI
VARLPPPIIPE
>TVAG_403280  | Trichomonas vaginalis G3 | Family T1, proteasome beta subunit, threonine peptidase | protein | length=191 beta4
MLSIVGLQGPDWVLIAADSSVSSSIICMSENYDRIAQLDDRHALAMSGETGDCLQLSEYL
QGNVALYKFRNGVELSSDALAHFIRHTMAKAVRKSPYEVNMLLSGYDGKPHLYFMDYLGT
LQSIPYGAQGYCQYFVMSVFDKHYKEGLTLEDGKELMKLALNQIKQRFTVAPHGFIVKLV
DKNGITKIDLE
>TVAG_290960  | Trichomonas vaginalis G3 | Family T1, proteasome beta subunit, threonine peptidase | protein | length=214 beta7
MQVITASGAIVAAKYDGGILLASDLSITYGSMFRHNNVSHFVEVAPNIIIGASGEFADFQ
TLIEVIKSVILQQQCKHNGEYLTASEVHNYIKRYMYQCRSNMKPLSCKVIVAGINPDGSK
FLACTDPYGASWESDHIGTGFGKYLQGLQIADVVNGSFDDVKKGITEVFRAVNARNTTAN
GKIEFITVTPQGINHLAPEQIDPNWEVVEGTWDQ
>TVAG_231360  | Trichomonas vaginalis G3 | Family T1, proteasome beta subunit, threonine peptidase | protein | length=275 beta2
MEAGLGFDFSNYARNKSLEPKLGKPQLLSTGTTIAAAIFDGGVVLGADTRATAGPIVAVK
DEMKLHYISDNIWVCGAGIAADNDNINAVISAKLRLFQMNTGLQPRVDQCTNILASRLFQ
YMGYIQAALIVGGIDFQGPQVYQVAPHGSFSKQPFIAQGSGSLAAISVLENRWHNKMNEH
DCMEMVADAIYAGITNDLGSGSHVNLCVIKRENPEDKQSKVIYTFYKDYRVPHENDRNFR
LEPQINNIDVEVIKTTERPLTLPDVHLEILDDAPA
>TVAG_103780  | Trichomonas vaginalis G3 | Family T1, proteasome alpha subunit, threonine peptidase | protein | length=232 alfa2
MGDSDFSLTTFSSGGKLNQIESALKAVSLGGQCVGVKAKNGAVIACESKPSSPLVEKVTN
LKVQKINDNVGIVYSGVNTDFHVILKSLRKASIKYSLRLGVEMPTREVVKHAAHKMQYYT
QIGGVRPFGVSLLIIGWEELGPTLWQVDPSGTFWAWKATALGKRSDGSRTFLERRYSEDQ
SVDDAIHTAISTLKEGFDGQLTAELIEIGVVDETRKFRTLSTAEIRDFLTEV
>TVAG_206040  | Trichomonas vaginalis G3 | Family T1, proteasome alpha subunit, threonine peptidase | protein | length=233 alfa6
MFRSKYDENATTFSPEGRILQVENAMKAVQQGMPTVGLKSKTHAVIAGVMHSPSEFSSHQ
PKIFKIDQHIGVAISGLTADGRGLCKFLRNECLHHTFCFGTEIRVADLADTVALQSQKKT
SKVGKRPYGVGLLMIGAGVDGPRLFETCPSGQHWEYNAQAIGRRAQAAKTYLETNLNEFP
DCTRDQLIRHALRALNDCKSRESDSLEAIALGVVGIDEPFTILEGPELQKYID
>TVAG_058050  | Trichomonas vaginalis G3 | Family T1, proteasome alpha subunit, threonine peptidase | protein | length=251 alfa3
MTYRYDAGTTTFSSDGRILQVEYAIQSINQAGTAIGVQFTNGVVLAAEKKNTGRLVDYLFPEKMAKIDGHIVTAVAGLTADANTLVDLMRTSAQKYLKTYDEQMPVEQLVRMVCDEKHSYTQYGGLRPYGVSFLIAGYDRHKGCQLYLTDPSGNFGGWKATAIGENNQTAQSILKSQYKDNMTATEAMDLTVKVLCKTLDSTSLSADKLEFAVLQFREEYGPKVRILTTSEVDTLMKRYEETIKKSAEEKE


Assembled  Tv20S proteasome calculated using Expasy (https://web.expasy.org/compute_pi/)
Theoretical pI/Mw: 5.90 / 348674.60 = 697,349 Da
```
